# Prevalence and mutational determinants of high tumor mutation burden in breast cancer

**DOI:** 10.1101/745265

**Authors:** R. Barroso-Sousa, E. Jain, O. Cohen, D. Kim, J. Buendia-Buendia, E. Winer, N. Lin, S.M. Tolaney, N. Wagle

**Affiliations:** Department of Medical Oncology, Dana-Farber Cancer Institute, Boston, MA, USA; Oncology Center, Hospital Sírio-Libanês Brasília, Brazil; Center for Cancer Precision Medicine, Dana-Farber Cancer Institute, Boston, MA, USA; Broad Institute of MIT and Harvard, Cambridge, MA, USA; Harvard Medical School, Boston, MA, USA; Department of Medicine, Brigham and Women’s Hospital, Boston, MA, USA

**Keywords:** breast cancer, tumor mutational burden, APOBEC, mutational signatures, immunotherapy, mismatch repair deficiency

## Abstract

**Background:** High tumor mutation burden (TMB) has been associated with benefit to immunotherapy in multiple tumor types. However, the prevalence of hypermutated breast cancer is not well described. The aim of this study is to evaluate frequency, mutational patterns, and genomic profile of hypermutated breast cancer.

**Patients and Methods:** We used de-identified data from individuals with primary or metastatic breast cancer from six different publicly available genomic studies. The prevalence of hypermutated breast cancer was determined among 3969 patients’ samples that underwent whole exome sequencing or gene panel sequencing. Samples were classified as having high TMB if they had ≥10 mutations per megabase (mut/Mb). An additional 8 patients were identified from a Dana-Farber Cancer Institute cohort for inclusion in the hypermutated cohort. Among patients with high TMB, the mutational patterns, and genomic profile were determined. A subset of patients was treated with regimens containing PD-1 inhibitors.

**Results:** The median TMB was 2.63 mut/Mb. Median TMB significantly varied according to tumor subtype (HR-/HER2-> HER2+ > HR+/HER2-, *p* < 0.05) and sample type (metastatic > primary, *p* 2.2×10^−16^). Hypermutated tumors were found in 198 patients (5%), with an enrichment in metastatic versus primary tumors (8.4% versus 2.9%, p = 6.5 × 10^−14^). APOBEC activity (59.2%), followed by mismatch repair deficiency (MMRd; 36.4%), were the most common mutational processes among hypermutated tumors. Three patients with hypermutated breast cancer—including two with a dominant APOBEC activity signature and one with a dominant MMRd signature—treated with pembrolizumab-based therapies derived an objective and durable response to therapy.

**Conclusion:** Hypermutation occurs in 5% of all breast cancers, with an enrichment in metastatic tumors. Different mutational signatures are present in this population, with APOBEC activity being the most common dominant process. Preliminary data suggest that hypermutated breast cancers are more likely to benefit from PD-1 inhibitors.

**Key Message:** High tumor mutation burden is found in 5% of all breast cancers and is more common in metastatic tumors. While different mutational signatures are present in hypermutated tumors, APOBEC activity is the most common dominant process. Preliminary data suggest that those tumors are more likely to benefit from PD-1 inhibitors.

## Introduction

Despite the success of immune checkpoint inhibitors (ICI) across several tumor types, to date, only a small fraction of patients with metastatic breast cancer (MBC) have shown benefit to PD-1/PD-L1 inhibitors given as monotherapy^1–6^. Thus, clinical trials have been launched to evaluate the efficacy of the combination of PD-1 axis inhibitors with other agents, including chemotherapy in breast cancer.

Recently, based on data from IMPASSION130, the US Food and Drug Administration (FDA) granted accelerated approval for the combination of atezolizumab plus nab-paclitaxel for the treatment of patients with metastatic triple-negative breast cancer (mTNBC) of tumors with 1% PD-L1 expression on immune cells in the tumor microenvironment^7^. However, other predictive biomarkers may help to increase the number of patients with breast cancer who are likely to benefit from ICI, including those with hormone receptor (HR)-positive disease.

It has been recognized that somatic mutations are the main source of tumor-specific antigens, or simply, neoantigens. Preclinical and clinical studies have shown that neoantigens are key targets of antitumor immunity.^8,9,10,11^ In this context, high tumor mutational burden (TMB) is associated with high neoantigen burden, high T-cell infiltration, and high response rates to immune checkpoint inhibitors across different tumor types.^12–21^ The objectives of this study are to evaluate the prevalence of hypermutation in breast tumors and determine the associated pathological characteristics, mutational signatures and genomic profiles. To do so, we analyzed publicly available genomic sequencing data from tumor samples from 3969 patients with breast cancer. We also present several patients with hypermutated breast cancer who were treated with PD-1/PD-L1 inhibitor-based regimens and achieved prolonged clinical benefit.

## Methods

### Patients and Samples

For the initial analysis, we evaluated de-identified genomic data from 3969 individuals with breast cancer from six different studies (Table 1). Whole exome sequencing (WES) data was obtained from The Cancer Genome Atlas breast cancer cohort (TCGA-BRCA) (http://gdac.broadinstitute.org/), The Metastatic Breast Cancer Project (MBCProject, April 2018) (https://www.mbcproject.org/data-release) and France study^22^, all publicly available on cbioportal.org (downloaded in May 2018). Gene panel sequencing data was obtained from the Dana-Farber Cancer Institute-OncoPanel (DFCI-OncoPanel), Memorial Sloan Kettering-Integrated Mutation Profiling of Actionable Cancer Targets (MSK-IMPACT) and Vanderbilt-Ingram Cancer Center (VICC), all found in the public release of AACR Project GENIE^23^, version 4.0, downloaded via Sage Synapse (http://synapse.org/genie). For individuals with multiple samples, sample with the highest TMB was chosen, hence using one sample per patient.

**Table 1 –.** Characteristics of different breast cancer datasets and frequency of hypermutation.

| Dataset | Patients | Type of tissue | Site of tissue |  |  | Genes sequenced (N) | Frequency of hypermutated tumors (%) |
| --- | --- | --- | --- | --- | --- | --- | --- |
|  |  |  | Primary Samples<br>(N (%)) | Metastatic Samples<br>(N (%)) | Unspecified or NA Samples<br>(N (%)) |  |  |
| France 2016<sup>+</sup> | 213 | FF | 0 (0.0) | 213 (100.0) | 0 (0.0) | ~20,000<sup>*</sup> | 4.2 |
| MBCProject April 2018<sup>#</sup> | 126 | FFPE | 100 (79.4) | 18 (14.3) | 8 (6.3) | ~20,000<sup>*</sup> | 3.2 |
| TCGA-BRCA Cell. 2015 | 977 | FF | 977 (100.0) | 0 (0.0) | 0 (0.0) | ~20,000<sup>*</sup> | 2.6 |
| GENIE-DFCI-ONCOPANEL-3 | 301 | FFPE | 176 (58.5) | 116 (38.5) | 9 (3.0) | 447 | 14.0 |
| GENIE-MSK IMPACT410 | 1009 | FFPE | 388 (38.5) | 621 (61.5) | 0 (0.0) | 410 | 7.3 |
| GENIE-MSK IMPACT468 | 1071 | FFPE | 696 (65.0) | 375 (35.0) | 0 (0.0) | 468 | 2.3 |
| GENIE-VICC-01-T5A | 92 | FFPE | 46 (50.0) | 46 (50.0) | 0 (0.0) | 322 | 7.6 |
| GENIE-VICC-01-T7 | 180 | FFPE | 72 (40.0) | 107 (59.4) | 1 (0.4) | 429 | 6.7 |
| **Total** | **3969** |  | **2455 (61.9)** | **1496 (37.7)** | **18 (0.04)** |  | **5.0** |

FF: Frozen; FFPE: Formalin fixed paraffin embedded; N: number.
<sup>#</sup>Provisional, April 2018; <sup>+</sup>Lefebvre et al. Plos Med 2016; <sup>\*</sup>Whole exome sequencing.

For the subsequent hypermutated cohort analysis, we also included 8 additional patients (from a cohort of 222 patients) from our ongoing study of estrogen receptor (ER)-positive metastatic breast cancer in the Center for Cancer Precision Medicine at Dana-Farber Cancer Institute (DFCI-CCPM).^24^ Prior to any study procedures, all patients provided written informed consent to whole exome sequencing of tumor and normal DNA, as approved by the Dana-Farber/Harvard Cancer Center Institutional Review Board (DF/HCC Protocol 05-246). Metastatic core biopsies were obtained from patients and samples were immediately snap frozen in optimal cutting temperature and stored in −80°C. Archived Formalin-Fixed Paraffin-Embedded (FFPE) blocks of primary tumor samples were also obtained.

### Assessment of TMB

TMB (mutation per megabase) was calculated as the total number of mutations detected for a given sample divided by the length of the total genomic target region captured with the exome or gene panel assay. The gene panels included had ≥ 1 Mb of target region captured. The TMB calculated from the specific gene panels selected for this analysis have previously been shown to have good correlation with TMB calculated from WES.^25–27^ The overall TMB distribution was used to identify the threshold for hypermutated tumors, using the following formula: median (TMB) + 2 × IQR(TMB), where IQR is the interquartile range. The calculated cutoff value was 9.44, which was rounded off to 10. Samples with TMB of ≥10 were classified as hypermutated.

#### Clinical annotations statistical analysis

TMB was correlated with available clinical annotations (sample type, receptor subtype and histology). These annotations were extracted from patient and tumor sample level clinical data from these studies. These annotations reflect the tumor characteristic at the time of tumor biopsy. Tumor biopsies from the TCGA-BRCA study were annotated as primary. In the France Study 2016, all tumor biopsies were designated as metastatic. In the MBCProject, tumor biopsies from the breast were designated as primary, except if there were clear clinical annotations that the breast biopsy was obtained in the metastatic setting, in which case they were designated as metastatic. Tumour biopsies from anatomic sites other than the breast in the MBCproject were designated as metastatic. For rest of the cohorts (obtained from AACR Project GENIE), tumor biopsies from the breast were designated as primary and biopsies from anatomic sites other than breast were designated as metastatic. Patients with complete clinical annotations were considered for statistical analysis. Wilcoxon test was used to calculate significance for differences in TMB across various clinical annotations. A p value of < 0.05 was considered to be statistically significant.

#### Immune cytolytic score calculation

Using RNA-sequencing data from the TCGA-BRCA dataset, we calculated the immune cytolytic activity defined as the geometric mean of expression values (RPKM) for the *GZMA* and *PRF1* genes.^12^

#### Neoantigen Prediction Analysis

Among the datasets mentioned above, we used the MBCProject for neoantigen prediction analysis since we had access to germline WES data. Using the Topiary tool (https://github.com/hammerlab/topiary), the mutated DNA sequences from WES were computationally translated into corresponding mutated peptide sequences. Patient specific human leukocyte antigen alleles were determined using Polysolver.^28^ NetMHC (v4.0)^29^ was used in order to predict MHC class I binding affinity for 8 to 11mer peptide sequences containing the mutated amino acid. Candidate neoantigens of mutated peptides were selected based on the following filters: binding affinity IC50 of ≤ 500nM to one (or more) of the patient-specific HLA alleles and percentile rank cutoff of ≤ 2.0.

#### Mutational Signature Analysis

Contributions of different mutation signatures were identified for each sample according to distribution of the six substitution classes (C>A, C>G, C>T, T>A, T>C, T>G) and the bases immediately 5′ and 3′ of the mutated base, producing 96 possible mutation subtypes. The extracted signatures were compared against the known and validated 30 COSMIC signatures.^30^ A sample was determined to have a dominant signature based on the maximum signature score attributable to that sample. We discuss four main signature categories here: homologous recombination deficiency-related (signature 3); activity of Apolipoprotein B mRNA Editing Catalytic Polypeptide-like (APOBEC) family (signatures 2 and 13); mismatch repair deficiency (MMRd; signatures 6, 15 and 20); and altered activity of POLE (signature 10). The analysis was performed using maftools package in R.^31^

#### Mutation Enrichment Analysis

Mutation rates for each gene and its differences were calculated for each patient. We restricted the analysis to known cancer driver genes as described in COSMIC Cancer Gene Consensus^32^ and PathCards^33^.

Fisher’s exact test was used to calculate significance. Multiple test correction was done using the p.adjust() function with false discovery rate method in R.

## Results

### Hypermutation across breast cancers

Genomic and clinical data from three WES studies (France Study 2016, MBCProject, and TCGA-BRCA) and three targeted panel studies (DFCI-ONCOPANEL, MSK-IMPACT and VICC) were combined to perform analysis on a total of 3969 patients with breast cancer. The frequency of hypermutation in breast cancers from each dataset (Figure 1A) varied from 2.3% (MSK-IMPACT468) to 14.0% (DFCI-OncoPanel). The median TMB across the entire cohort was 2.63, with a range of 0.2–290.8 (Figure 1B). Overall, 5% (198 cases) of breast cancers analyzed were hypermutated based on the calculated cutoff of 10 mutations/megabase (see methods).

**Figure 1 –.**
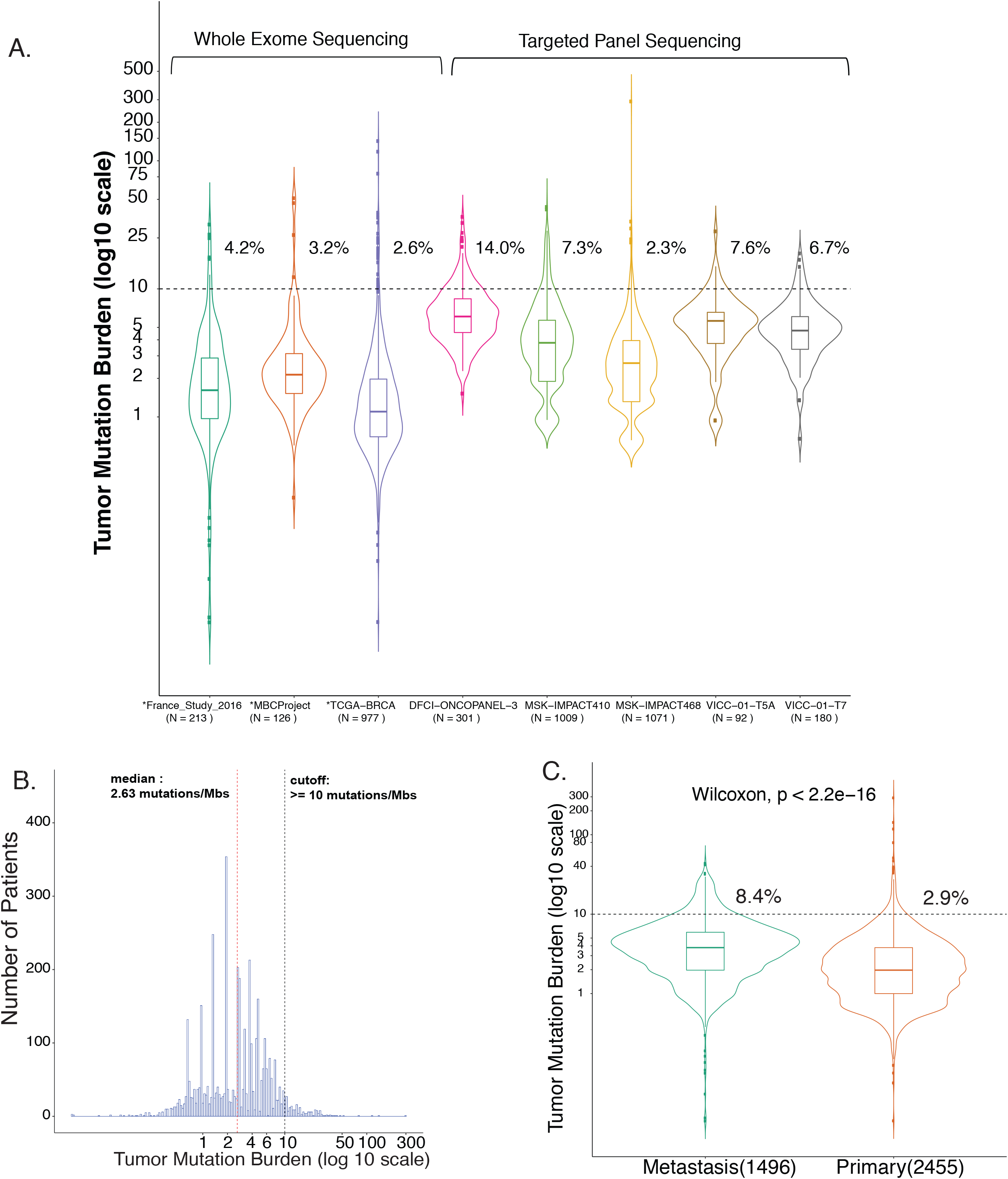
Tumor Mutation Burden across 3969 primary and metastatic breast cancers. (A) TMB (y-axis) distribution for each dataset (x-axis) used in the analysis. Sample points above the black dotted line at 10 mutations/megabase represents the hypermutated tumors. Percentage of hypermutation is indicated for each dataset. Datasets marked with * on x-axis used whole exome sequencing; the remaining datasets used several-hundred gene targeted sequencing panels. Numbers in parentheses represents the total number of patients included in this analysis from each dataset. (B) Histogram indicating the mutation burden across 3969 samples. Red dotted line indicates median TMB and black dotted line indicates the cutoff chosen to define hypermutation. (C) Boxplot representing median TMB for metastatic tumors versus primary tumors. Abbreviations: TMB: tumor mutational burden.

Metastatic tumors (see Methods), had a higher median TMB compared to primary tumors (3.8 vs 2.0, p <2.2 × 10^−16^) (Figure 1C). There was no significant correlation between TMB and age at diagnosis (R^2^ = 0.13, p = 3.6×10^−5^) (Supplementary Figure 1A) and no significant difference in TMB across histology types (p = 0.074) (Supplementary Figure 1B). Triple-negative breast cancers (TNBC) had significantly higher median TMB (1.8) compared to HR-positive cancers (1.1, p = 2.8 × 10^−8^) or HER2-positive cancers (1.3, p = 0.003) (Supplementary Figure 1C).

Among hypermutated cases, the median TMB was 14.46. We analyzed the hypermutated tumors according to clinical and pathological characteristics (Supplementary Table 1). The frequency of hypermutated breast cancer was higher for metastatic samples as compared to primary samples (8.4% vs 2.9%, Fisher’s exact, p = 6.529 × 10^−14^). Additionally, 8.7% of invasive lobular carcinomas were hypermutated as compared to 4.0% of invasive ductal carcinomas, but this difference in prevalence was not statistically significant (p = 0.074). However, among the metastatic samples, we observed a significant enrichment of hypermutation in metastatic ILC (17.0%) as compared to metastatic IDC tumors (7.8%) (Fisher’s exact, p= 0.001782). The prevalence of hypermutated breast cancer was similar among different disease receptor subtypes (3.7-3.9%).

To compare the differences in mutation rate between distant metastatic biopsies, primary biopsies from patients who eventually developed metastases, and primary biopsies from patients who did not develop metastases, we performed several exploratory comparisons, though the numbers of samples used for these comparisons was small. The frequency of hypermutation in distant metastatic tumors biopsies (France Study 2016 and MBCProject; 4 out of 123) was similar when compared to primary tumors (TCGA-BRCA, 25 out of 977)(3.1% vs 2.6%, Fisher’s exact, p=0.7). The frequency of hypermutation among primary tumors which eventually became metastatic (MBCProject; 3 out of 126) was similar to that of primary tumors overall, most of which did not recur (TCGA-BRCA,N=25), (2.4% vs 2.6%, Fisher’s exact p=1).

### Hypermutated breast cancers have a higher cytolytic score and higher neoantigen burden

We evaluated whether hypermutation correlates with an increased immune cytolytic activity, which has been used as a surrogate of tumor-infiltrating lymphocytes (TILs)^12^. Using RNA sequencing data available in the TCGA-BRCA dataset (N =974), we observed that hypermutated breast tumors (N = 25) had higher cytolytic activity compared to non-hypermutated breast tumors (p 0.0048; Supplementary Figure 2A).

Neoantigen burden was evaluated in tumor samples from the MBCProject (N = 157), in which we were able to analyze both germline and tumor WES data. The 4 hypermutated breast cancers in this cohort had a higher neoantigen burden compared to the 153 non-hypermutated tumors (Supplementary Figure 2B). Together. this data suggests that hypermutated breast cancers may have increased T-cell infiltration.

### APOBEC is the dominant mutational process among hypermutated breast cancers

We next investigated the potential drivers of hypermutation in breast cancer by assessing the mutational signatures present in these hypermutated tumors (Figure 2). Mutational processes causing cancer can arise due to intrinsic dysfunction (defective DNA replication, enzymatic modification of DNA and defective DNA repair) or extrinsic factors (exposure to ultraviolet light, mutagens or tobacco smoke). These mutational processes generate unique patterns of mutation types, which are termed as mutational signatures. We found that most hypermutated breast cancers (59.2%) have a dominant APOBEC activity signature (signature 2 and 13). APOBEC signature has been attributed to the activity of the AID/APOBEC family of cytidine deaminases converting cytosine to uracil. When dysregulated, this family of enzymes can be a major source of mutations in several cancers, including non-small cell lung cancer and bladder cancer.^34^

**Figure 2.**
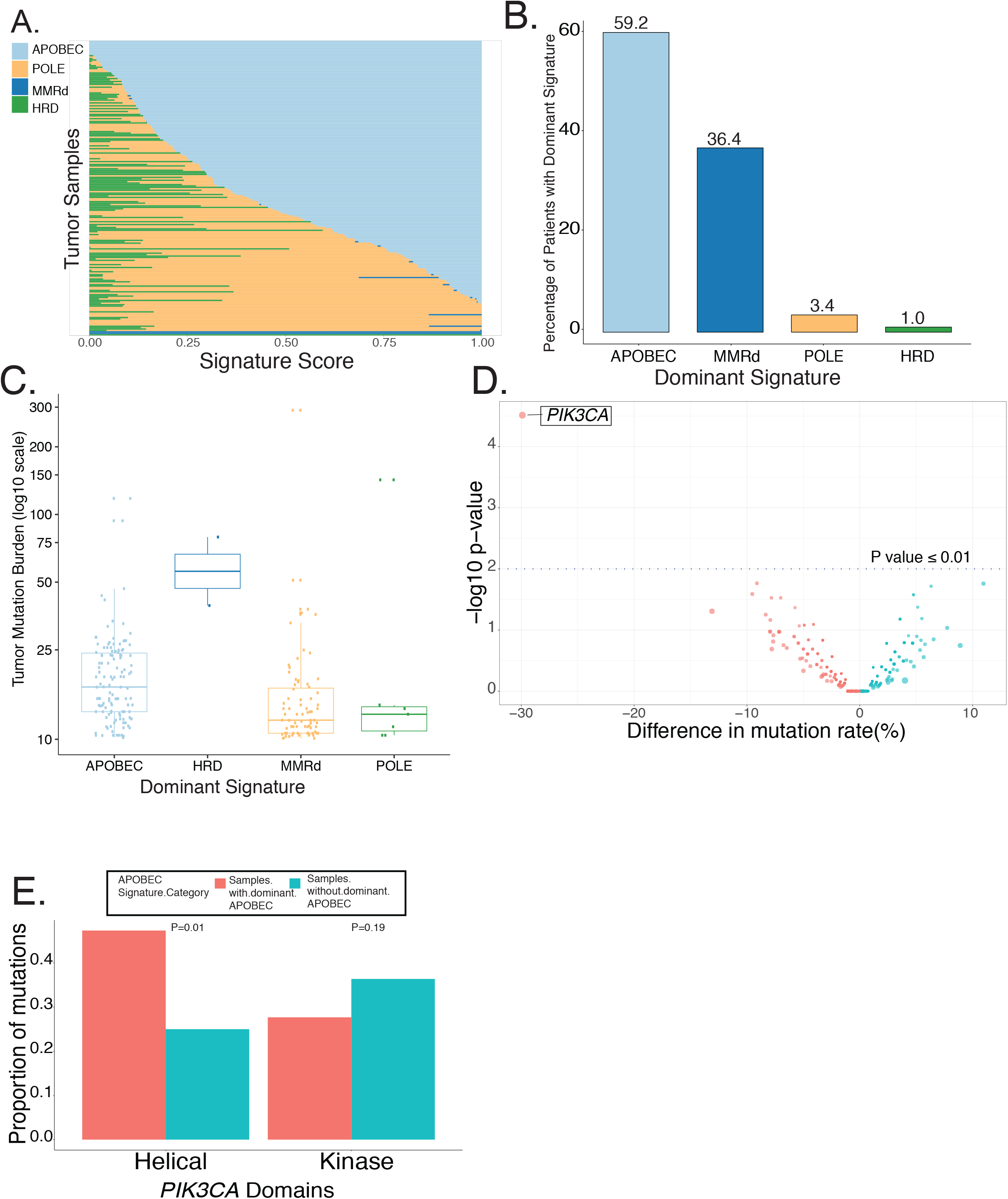
Mutational signatures prevalent in hypermutated breast cancer. (A) Signature score proportions (x-axis) for each of the 206 hypermutated patients (y-axis). Light blue represents APOBEC activity; blue represents DNA mismatch repair deficiency; light orange represents homologous recombination deficiency; green represents POLE signature (B) Each bar represents the percentage of patients across four dominant signatures sub groups. 59.2% of patients have dominant APOBEC signature. (C) TMB (y-axis log scale) distribution across four dominant signatures detected. (D) Volcano plot indicating mutational rate differences (x-axis) for each gene (represented as a dot). Red colored dot are genes having higher mutation rate in dominant APOBEC hymer-mutated tumors. Green colored dot are genes having higher mutation rate in non-dominant APOBEC hymermutated tumors. *PIK3CA* is significantly enriched in the APOBEC high hymermutated tumors. Y-axis represents negative log scale of P value. (E) *PIK3CA* alteration proportions for helical and kinase domain in dominant APOBEC hypermutated tumors (red) and non-dominant APOBEC hypermutated tumors (blue).

In another 36.4% of the hypermutated breast cancers, we found dominant signature signifying mismatch repair deficiency (MMRd) (signature 6, 15 and 20). MMRd leads to hypermutation and this signature is associated with high numbers of small insertions and deletions at mono/polynucleotide repeats regions.

Other patients exhibited different dominant mutational signatures: ~1.0% were found with a signature of homologous recombination deficiency (signature 3). Signature 3 is known to be strongly associated with *BRCA1/2* mutations in breast cancer. This signature is characterized by the presence of larger deletions with overlapping microhomology at breakpoint junctions. Furthermore, 3.4% patients presented with a dominant signature associated with the altered activity of error prone DNA polymerase epsilon (POLE) (signature 10) (Figure 2A and 2B). Signature 10 is known to cause ultra-hypermutation in small proportion of tumors in colorectal and uterine cancer.^30^ The median TMB was higher for samples with dominant APOBEC and homologous recombination deficiency signatures (17.1 and 59.4, respectively), followed by tumors with dominant POLE and MMRd signatures (12.2 and 12.9, respectively, Figure 2C).

### Genomic landscape of APOBEC high and low hypermutated breast cancers

Given the high proportion of hypermutated breast cancers with dominant APOBEC signatures, we sought to determine if there were any differences in the genomic landscape between hypermutated tumors that had a dominant APOBEC signature versus hypermutated tumors without a dominant APOBEC signature (Figure 2D). *PIK3CA* was found to be mutated in 68.6% of hypermutated tumors with dominant APOBEC signature versus 37.6% of hypermutated tumors without dominant APOBEC signature (p 1.63×10^−5^, q 0.015). The proportion of *PIK3CA* mutations in the helical domain were enriched within hypermutated tumors with dominant APOBEC signature (47.3% vs 25.0%; p 0.01; Figure 2E) and mutations in the kinase domain were enriched in hypermutated tumors without a dominant APOBEC signature (p 0.19). Detailed *PIK3CA* alterations counts in different gene domains in hypermutated tumors are presented in Supplementary Table 2.

### Response to anti-PD-1/PD-L1 based therapies in hypermutated breast cancer

Prior studies have demonstrated a correlation between hypermutation and response to immune checkpoint inhibitors.^35–37^ However, at present immune checkpoint inhibitors are only approved for breast cancers with MMR deficiency. We hypothesized that hypermutated breast cancers may respond to immune checkpoint inhibitors regardless of the underlying mutational signature. To test this, we examined the treatment histories of 222 patients with metastatic breast cancer from our prospective metastatic biopsy cohort at DFCI^24^. We identified 8 pts (3.6%) with hypermutated breast cancer, of whom four had received treatment with anti-PD-1/PD-L1 based therapies. Notably, three of these patients achieved an objective response to therapy and prolonged progression-free interval (Figure 3). Detailed prior systemic treatment details for these three patients are presented in Supplementary Table 3. Response to therapy was not able to be assessed in the fourth patient who received immune checkpoint inhibitor therapy, as this patient had central nervous system metastasis, including leptomeningeal disease, and died two weeks after starting therapy with an anti-PD-1 antibody. Analysis of mutational signatures in the metastatic biopsies demonstrated dominant APOBEC activity signatures in two patients and a dominant MMRd signature in the third one.

**Figure 3.**
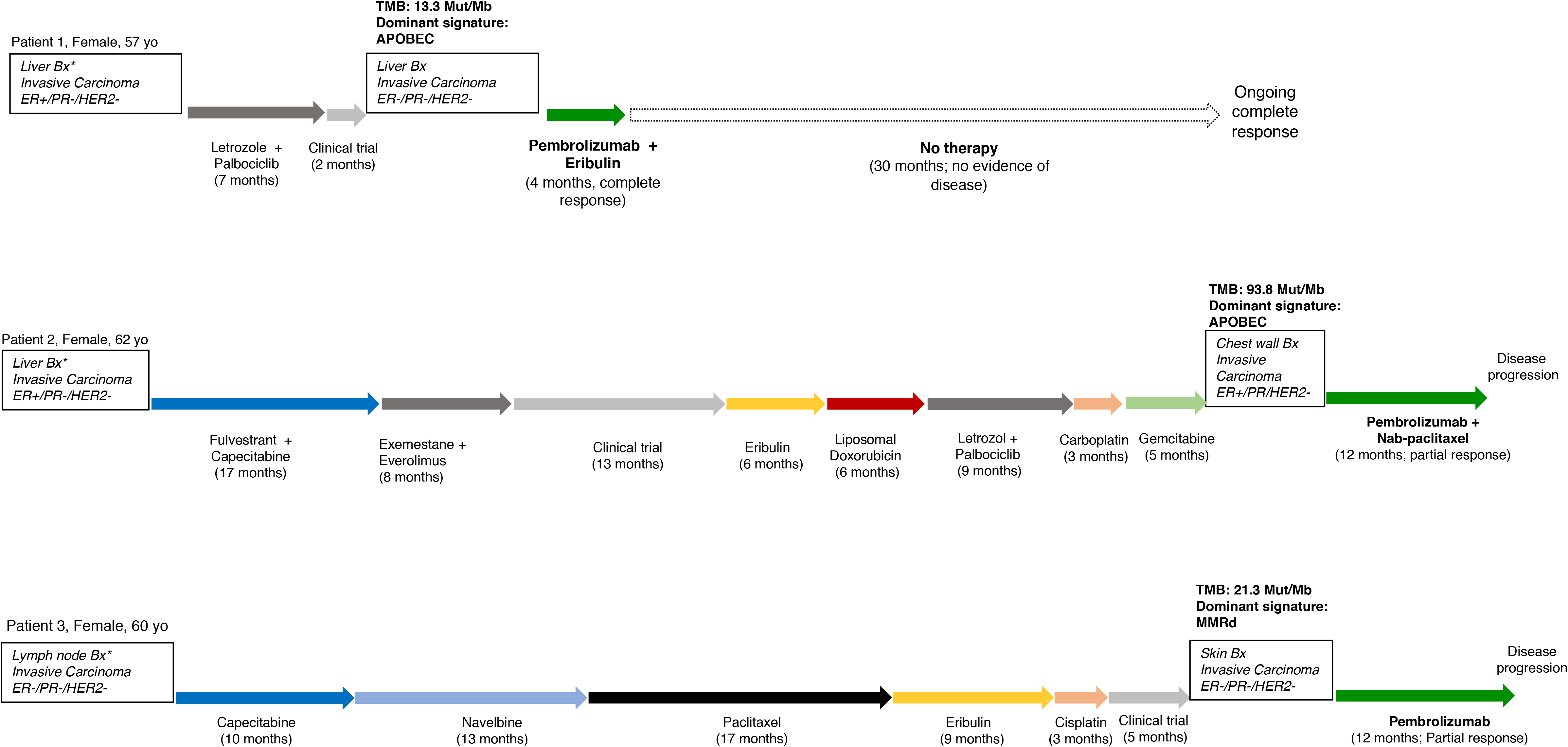
Details of the treatment received in the metastatic setting of patients with advanced breast cancer included in DFCI biobank cohort and treated with pembrolizumab-based therapy. APOBEC: Apolipoprotein B mRNA-editing enzyme catalytic polypeptide-like 3; Bx: biopsy; Dx: diagnosis; ER: estrogen receptor; IHC: immunohistochemistry; MMRd: mismatch repair deficiency; PR progesterone receptor; TMB: tumor mutational burden.

## Discussion

To our knowledge, this work represents the largest study evaluating the prevalence and the mutational drivers of hypermutated breast cancers. Using large scale sequencing data from six different breast cancer cohorts with a total of 3969 patients, we found that the prevalence of hypermutation in breast tumors was 5%. Notably, the prevalence of hypermutation was significantly higher in metastatic tumors than in primary tumors (8.4% vs 2.9%, Fisher’s exact, p = 6.529 × 10^−14^) and particularly in metastatic ILC versus metastatic IDC (17.0% versus 7.8%, Fisher’s exact, p= 0.001782). Analysis of a subset of these tumors demonstrated that hypermutated tumors had a higher neoantigen burden and a higher cytolytic score compared to non-hypermutated tumors. APOBEC activity was found to be the most common dominant mutational process associated with hypermutation in breast cancer, present in more than half of the hypermutated tumors. A dominant MMRd signature was present in an additional 36% of hypermutated breast cancers. Finally, we presented the histories of three patients with hypermutated breast cancer – 2 with a dominant APOBEC activity signature and 1 with a dominant MMRd signature – who achieved objective and durable responses following pembrolizumab-based regimens.

Our study found that 5% of patients had hypermutated tumors with 8.4% of metastatic lesions being hypermutated versus 2.9% of primary tumors. It is not clear why there is an enrichment of hypermutated breast tumors in metastatic samples. It is possible that this hypermutated phenotype is acquired during tumor evolution and could be associated with resistance to prior systemic therapy or with the development of metastases. In fact, mutational signature 13 that is associated with APOBEC activity and hypermutation is observed late in breast cancer evolution.^38–40^ Another possibility could be that hypermutation is also enriched in primaries that ultimately become metastatic and thus may be a poor prognostic factor. However, using an exploratory analysis in a subset of samples, we did not find significant enrichment of hypermutation in primary tumors of patients which ultimately became metastatic (MBCProject) when compared to primary tumors in general, most of which did not recur (TCGA-BRCA) (2.4% vs 2.6%, Fisher’s exact test, p = 1). The frequency of hypermutation was also similar in distant metastatic biopsies (France Study 2016 and MBCProject) when compared to primary tumors (TCGA-BRCA) (3.1 % vs 2.6%, Fisher’s exact test p = 0.7). Additional studies with sufficient sample size evaluating hypermutated metastatic biopsies paired with corresponding primaries from the same individuals will be necessary to further clarify this issue.

Data from IMPASSION130 established PD-L1 expression on immune cells as a predictive biomarker of benefit to atezolizumab plus chemotherapy in mTNBC^7^. However, there are still controversies surrounding the broad utility of PD-L1 expression for selecting patients for immunotherapy. Some of the concerns include that fact that PD-L1 is a dynamic marker, with varying expression over time. Additionally, data suggests discordance amongst pathologists in determining PD-L1 positivity. Perhaps more importantly some patients who test positive for PD-L1 may not respond to the therapy, and some patients who test negative may still respond^41^. Altogether, this has led to the investigation of additional biomarkers to predict benefit or resistance to immunotherapy. Across different tumor types, high TMB has been associated with improved clinical benefit to ICI.^14,15,19,20,35–37^ Notably, it has been shown that TMB and PD-L1 expression are independently predictive markers of response to ICI and have low correlation across multiple tumors^35^.

A better understanding of the forces driving hypermutation in breast cancers may be clinically relevant. The FDA granted accelerated approval for pembrolizumab in any MMRd tumors, and for the combination of nivolumab plus ipilimumab to treat refractory MMRd colorectal cancers. Given that MMR defects are one of the most important mechanisms associated with hypermutation, we investigated whether these hypermutated breast tumors were also associated with MMRd. Our study showed that while 36.4% of these hypermutated tumors have a dominant MMRd signature, the vast majority (59.2%) presented with a dominant APOBEC mutational signature. Importantly, this suggests that most hypermutated breast tumors will be missed if we only search for markers of MMRd or microsatellite instability.

Further studies should be done to confirm whether high TMB is predictive of benefit to immunotherapy in solid tumors independent of the mutational driver. While this is not yet known, in non-small cell lung cancer, APOBEC mutational signature was shown to be specifically enriched in patients with durable clinical benefit after immunotherapy.^40^ Additionally, APOBEC upregulation correlates with high levels of PD-L1 expression.^42^ Recently, Goodman *et al*. suggested that *PDL1* amplification correlates with improved responses to ICI.^43^ Therefore, it is conceivable that such genomic alteration works as a mechanism of immune escape from an endogenous immune response in tumors with APOBEC dysregulation. In addition, and in agreement with other studies,^44,45^ we found a relationship between APOBEC-induced mutagenesis and *PIK3CA* mutations, especially with mutations in the helical domain (Figure 2E). Miao *et al*. reported that *PIK3CA* mutations were associated with complete or partial response to immune checkpoint therapy in microsatellite-stable solid tumors.^46^

Notably, in our DFCI metastatic HR-positive breast cancer cohort, four patients with hypermutated tumors have received ICI-based therapies. Three patients achieved objective and durable responses: one received pembrolizumab given as monotherapy as part of the trial NCT02447003; one received the combination of pembrolizumab plus eribulin as part of the NCT02513472; and one received pembrolizumab plus nab-paclitaxel outside of a clinical trial. Interestingly, two of them had dominant APOBEC activity signatures while the other had a dominant MMRd signature. To better evaluate whether hypermutated breast cancers are responsive to immunotherapy, we launched a multicenter, single arm trial (NIMBUS), phase II trial of nivolumab plus ipilimumab in metastatic hypermutated HER2-negative breast cancers (NCT03789110). Patients are eligible for this trial if they have metastatic HER2-negative breast cancer with >= 10 mut/Mb as assessed by larger targeted panels (>300 genes) and have not been treated with more than three prior lines of systemic therapy in the metastatic setting.

Given that more than 250,000 women and men are diagnosed with breast cancer in the U.S. every year, a frequency of 5% means that over 13,000 patients with hypermutated breast cancer being diagnosed annually just in the United States. This number suggests that hypermutated breast cancers are more prevalent other cancer subtypes such as non-small cell lung cancers with ALK-rearrangements or *ROS1* translocation, in which targeted therapy is successfully applied. Furthermore, enrichment in the frequency of hypermutation among metastatic ILC is notable and brings up the question whether all ILC should be investigated for hypermutation.

### Strengths and Limitations

The strengths of this study include the large sample size, the inclusion of subsets of patients with WES and RNA sequencing, the substantial number of patients with metastatic biopsies, and the mutational signature analysis. However, our study has some limitations. First, clinical annotation data was unavailable in some studies, especially regarding receptor subtypes. There might be differences in definition of clinical annotations (e.g. metastasis vs primary) across different cohorts. While GENIE study defines metastasis vs. primary based on the site of acquisition of the tumor tissue, studies like the MBCProject define it based on the stage of disease (primary or metastatic) when the tumor tissue was acquired. Second, we performed a combined analysis of different datasets and batch effect and cohort bias are possible. Although previous studies have indicated a high concordance between findings of similar studies using different technological tools^47^, we acknowledge the caveats of comparing TMB using different platforms. TMB is influenced by tumor purity, ploidy, sequencing depth of coverage, and analysis methodologies. Since we are using publicly available data that has already been analyzed, we were not able to reanalyze and recalculate TMB it for each of the data sets in a standardized manner.

In addition, the definition of high TMB is still not optimized across cancer subtypes^19,48^, including breast cancer. The cutoff used to define hypermutation is consistent with the one used in large pan-cancer analysis conducted by Campbell *et al*.^25,26^ Our study used a combination of targeted gene panel and WES to determine the TMB cutoff, majority of samples coming from targeted gene panels. Although gene panels tend to estimate higher mutation burden, we selected larger gene panels which are known to have good correlation with WES with respect to TMB calculation.^25–27^ There are multiple ongoing initiatives to standardize TMB assessment, and further work is necessary to establish the best cutoff for using TMB as a predictive biomarker of response to immunotherapy.^49,50^

## Conclusion

Our data suggest 5% of breast cancer have a high TMB, with an enrichment in metastatic tumors. These tumors are associated with a higher neoantigen burden and are more T-cell infiltrated. Furthermore, different mutational signatures are present in this population, with APOBEC activity being the most common dominant mutational process. Preliminary data suggest that hypermutated breast cancers are more likely to benefit from ICI supporting the conduct of the ongoing NIMBUS trial.

## Acknowledgements

We thank Karla Helvie, Laura Dellostritto, Lori Marini, Nelly Oliver, Shreevidya Periyasamy, Colin Mackichan, and Max Lloyd for assistance with the DFCI patient sample collection and annotation. We thank Dr. Elizabeth Mittendorf for helpful discussions and comments on the manuscript. We thank Kaitlyn Bifolck for her editorial support to this manuscript. We are grateful to all the patients who volunteered for research protocols and generously provided the tissue analyzed in this study.

## References

1. Nanda R, Chow LQM, Dees EC, et al. Pembrolizumab in Patients With Advanced Triple-Negative Breast Cancer: Phase Ib KEYNOTE-012 Study. J Clin Oncol. 2016;34(21):2460–2467.

2. Adams S, Loi S, Toppmeyer D, et al. Phase 2 study of pembrolizumab as first-line therapy for PD-L1-positive metastatic triple-negative breast cancer (mTNBC): Preliminary data from KEYNOTE-086 cohort B. Journal of Clinical Oncology. 2017;35(15_suppl): 1088–1088. doi: 10.1200/jco.2017.35.15_suppl.1088

3. Adams S, Schmid P, Rugo HS, et al. Phase 2 study of pembrolizumab (pembro) monotherapy for previously treated metastatic triple-negative breast cancer (mTNBC): KEYNOTE-086 cohort A. Journal of Clinical Oncology. 2017;35(15_suppl):1008–1008. doi: 10.1200/jco.2017.35.15_suppl.1008

4. Loi S, Giobbe-Hurder A, Gombos A, et al. Abstract GS2-06: Phase Ib/II study evaluating safety and efficacy of pembrolizumab and trastuzumab in patients with trastuzumab-resistant HER2-positive metastatic breast cancer: Results from the PANACEA (IBCSG 45-13/BIG 4-13/KEYNOTE-014) study. Cancer Research. 2018;78(4 Supplement):GS2–GS06. doi:10.1158/1538-7445.sabcs17-gs2-06

5. Dirix LY, Takacs I, Jerusalem G, et al. Avelumab, an anti-PD-L1 antibody, in patients with locally advanced or metastatic breast cancer: a phase 1b JAVELIN Solid Tumor study. Breast Cancer Res Treat. 2018;167(3):671–686.

6. Rugo HS, Delord J-P, Im S-A, et al. Safety and Antitumor Activity of Pembrolizumab in Patients with Estrogen Receptor–Positive/Human Epidermal Growth Factor Receptor 2–Negative Advanced Breast Cancer. Clinical Cancer Research. 2018;24(12):2804–2811. doi:10.1158/1078-0432.ccr-17-3452

7. Schmid P, Adams S, Rugo HS, et al. Atezolizumab and Nab-Paclitaxel in Advanced Triple-Negative Breast Cancer. New England Journal of Medicine. 2018;379(22):2108–2121. doi: 10.1056/nejmoa1809615

8. Matsushita H, Vesely MD, Koboldt DC, et al. Cancer exome analysis reveals a T-cell-dependent mechanism of cancer immunoediting. Nature. 2012;482(7385):400–404. doi:10.1038/nature10755

9. DuPage M, Mazumdar C, Schmidt LM, Cheung AF, Jacks T. Expression of tumour-specific antigens underlies cancer immunoediting. Nature. 2012;482(7385):405–409. doi:10.1038/nature10803

10. Brown SD, Warren RL, Gibb EA, et al. Neo-antigens predicted by tumor genome meta-analysis correlate with increased patient survival. Genome Research. 2014;24(5):743–750. doi: 10.1101/gr.165985.113

11. Kreiter S, Vormehr M, van de Roemer N, et al. Mutant MHC class II epitopes drive therapeutic immune responses to cancer. Nature. 2015;520(7549):692–696. doi:10.1038/nature14426

12. Rooney MS, Shukla SA, Wu CJ, Getz G, Hacohen N. Molecular and genetic properties of tumors associated with local immune cytolytic activity. Cell. 2015;160(1-2):48–61.

13. Giannakis M, Mu XJ, Shukla SA, et al. Genomic Correlates of Immune-Cell Infiltrates in Colorectal Carcinoma. Cell Rep. 2016;17(4):1206.

14. Snyder A, Makarov V, Merghoub T, et al. Genetic basis for clinical response to CTLA-4 blockade in melanoma. N Engl J Med. 2014;371(23):2189–2199.

15. Rizvi NA, Hellmann MD, Snyder A, et al. Cancer immunology. Mutational landscape determines sensitivity to PD-1 blockade in non-small cell lung cancer. Science. 2015;348(6230): 124–128.

16. Van Allen EM, Miao D, Schilling B, et al. Genomic correlates of response to CTLA-4 blockade in metastatic melanoma. Science. 2015;350(6257):207–211.

17. Le DT, Uram JN, Wang H, et al. PD-1 Blockade in Tumors with Mismatch-Repair Deficiency. New England Journal of Medicine. 2015;372(26):2509–2520. doi:10.1056/nejmoa1500596

18. Carbone DP, Reck M, Paz-Ares L, et al. First-Line Nivolumab in Stage IV or Recurrent Non-Small-Cell Lung Cancer. N Engl J Med. 2017;376(25):2415–2426.

19. Hellmann MD, Ciuleanu T-E, Pluzanski A, et al. Nivolumab plus Ipilimumab in Lung Cancer with a High Tumor Mutational Burden. N Engl J Med. 2018;378(22):2093–2104.

20. Hellmann MD, Callahan MK, Awad MM, et al. Tumor Mutational Burden and Efficacy of Nivolumab Monotherapy and in Combination with Ipilimumab in Small-Cell Lung Cancer. Cancer Cell. 2018;33(5):853–861.e4. doi:10.1016/j.ccell.2018.04.001

21. Campesato LF, Barroso-Sousa R, Jimenez L, et al. Comprehensive cancer-gene panels can be used to estimate mutational load and predict clinical benefit to PD-1 blockade in clinical practice. Oncotarget. 2015;6(33):34221–34227.

22. Lefebvre C, Bachelot T, Filleron T, et al. Mutational Profile of Metastatic Breast Cancers: A Retrospective Analysis. PLoS Med. 2016;13(12):e1002201.

23. AACR Project GENIE Consortium. AACR Project GENIE: Powering Precision Medicine through an International Consortium. Cancer Discov. 2017;7(8):818–831.

24. Cohen O, Kim D, Oh C, et al. Abstract S1-01: Whole exome and transcriptome sequencing of resistant ER metastatic breast cancer. Cancer Research. 2017;77(4 Supplement):S1–01. doi: 10.1158/1538-7445.sabcs16-s1-01

25. Campbell BB, Light N, Fabrizio D, et al. Comprehensive Analysis of Hypermutation in Human Cancer. Cell. 2017;171(5):1042–1056.e10. doi:10.1016/j.cell.2017.09.048

26. Garofalo A, Sholl L, Reardon B, et al. The impact of tumor profiling approaches and genomic data strategies for cancer precision medicine. Genome Med. 2016;8(1):79.

27. Rizvi H, Sanchez-Vega F, La K, et al. Molecular Determinants of Response to Anti-Programmed Cell Death (PD)-1 and Anti-Programmed Death-Ligand 1 (PD-L1) Blockade in Patients With Non-Small-Cell Lung Cancer Profiled With Targeted Next-Generation Sequencing. J Clin Oncol. 2018;36(7):633–641.

28. Shukla SA, Rooney MS, Rajasagi M, et al. Comprehensive analysis of cancer-associated somatic mutations in class I HLA genes. Nature Biotechnology. 2015;33(11): 1152–1158. doi: 10.1038/nbt.3344

29. Andreatta M, Nielsen M. Gapped sequence alignment using artificial neural networks: application to the MHC class I system. Bioinformatics. 2016;32(4):511–517. doi: 10.1093/bioinformatics/btv639

30. Alexandrov LB, Nik-Zainal S, Wedge DC, Campbell PJ, Stratton MR. Deciphering Signatures of Mutational Processes Operative in Human Cancer. Cell Reports. 2013;3(1):246–259. doi: 10.1016/j.celrep.2012.12.008

31. Mayakonda A, Lin D-C, Assenov Y, Plass C, Koeffler HP. Maftools: efficient and comprehensive analysis of somatic variants in cancer. Genome Res. 2018;28(11): 1747–1756.

32. Sondka Z, Bamford S, Cole CG, Ward SA, Dunham I, Forbes SA. The COSMIC Cancer Gene Census: describing genetic dysfunction across all human cancers. Nat Rev Cancer. 2018;18(11):696–705.

33. Belinky F, Nativ N, Stelzer G, et al. PathCards: multi-source consolidation of human biological pathways. Database. 2015;2015. doi:10.1093/database/bav006

34. Roberts SA, Lawrence MS, Klimczak LJ, et al. An APOBEC cytidine deaminase mutagenesis pattern is widespread in human cancers. Nat Genet. 2013;45(9):970–976.

35. Cristescu R, Mogg R, Ayers M, et al. Pan-tumor genomic biomarkers for PD-1 checkpoint blockade-based immunotherapy. Science. 2018;362(6411). doi:10.1126/science.aar3593

36. Ott PA, Bang Y-J, Piha-Paul SA, et al. T-Cell-Inflamed Gene-Expression Profile, Programmed Death Ligand 1 Expression, and Tumor Mutational Burden Predict Efficacy in Patients Treated With Pembrolizumab Across 20 Cancers: KEYNOTE-028. J Clin Oncol. 2019;37(4):318–327.

37. Samstein RM, Lee C-H, Shoushtari AN, et al. Tumor mutational load predicts survival after immunotherapy across multiple cancer types. Nat Genet. 2019;51(2):202–206.

38. Nik-Zainal S, Van Loo P, Wedge DC, et al. The life history of 21 breast cancers. Cell. 2012;149(5): 994–1007.

39. Nik-Zainal S, Davies H, Staaf J, et al. Landscape of somatic mutations in 560 breast cancer whole-genome sequences. Nature. 2016;534(7605):47–54.

40. Wang S, Jia M, He Z, Liu X-S. APOBEC3B and APOBEC mutational signature as potential predictive markers for immunotherapy response in non-small cell lung cancer. Oncogene. 2018;37(29):3924–3936.

41. Ribas A, Hu-Lieskovan S. What does PD-L1 positive or negative mean? J Exp Med. 2016;213(13):2835–2840.

42. Boichard A, Tsigelny IF, Kurzrock R. High expression of PD-1 ligands is associated with mutational signature and APOBEC3 alterations. Oncoimmunology. 2017;6(3):e1284719.

43. Goodman AM, Piccioni D, Kato S, et al. Prevalence of PDL1 Amplification and Preliminary Response to Immune Checkpoint Blockade in Solid Tumors. JAMA Oncol. 2018;4(9): 1237–1244.

44. McGranahan N, Favero F, de Bruin EC, Birkbak NJ, Szallasi Z, Swanton C. Clonal status of actionable driver events and the timing of mutational processes in cancer evolution. Science Translational Medicine. 2015;7(283):283ra54–ra283ra54. doi: 10.1126/scitranslmed.aaa1408

45. Temko D, Tomlinson IPM, Severini S, Schuster-Böckler B, Graham TA. The effects of mutational processes and selection on driver mutations across cancer types. Nat Commun. 2018;9(1): 1857.

46. Miao D, Margolis CA, Vokes NI, et al. Genomic correlates of response to immune checkpoint blockade in microsatellite-stable solid tumors. Nat Genet. 2018;50(9): 1271–1281.

47. Van Allen EM, Robinson D, Morrissey C, et al. A comparative assessment of clinical whole exome and transcriptome profiling across sequencing centers: implications for precision cancer medicine. Oncotarget. 2016;7(33). doi:10.18632/oncotarget.9184

48. Legrand FA, Gandara DR, Mariathasan S, et al. Association of high tissue TMB and atezolizumab efficacy across multiple tumor types. Journal of Clinical Oncology. 2018;36(15_suppl): 12000–12000. doi: 10.1200/jco.2018.36.15_suppl.12000

49. Stenzinger A, Allen JD, Maas J, et al. Tumor mutational burden standardization initiatives: Recommendations for consistent tumor mutational burden assessment in clinical samples to guide immunotherapy treatment decisions. Genes, Chromosomes and Cancer. 2019. doi: 10.1002/gcc.22733

50. Chan TA, Yarchoan M, Jaffee E, et al. Development of tumor mutation burden as an immunotherapy biomarker: utility for the oncology clinic. Annals of Oncology. 2019;30(1):44–56. doi: 10.1093/annonc/mdy495

