## Supplemental Tables and Figures for "Prevalence and mutational determinants of high tumor mutation burden in breast cancer"

Supplemental Tables: 3

Supplemental Figures: 2

**Supplementary Table 1**- Clinical characteristics of 198 (5%) hypermutated patient identified from seven different studies.

| | Total Number of Patients | Number of Hypermutated Patients | Percentage of Hypermutated Patients (%) |
|---|---|---|---|
| SAMPLE TYPE | | | |
| METASTASIS | 1371 | 125 | 8.4 |
| PRIMARY | 2384 | 71 | 2.9 |
| UNSPECIFIED | 9 | 1 | 10.0 |
| NA | 7 | 1 | 12.5 |
| RECEPTOR STATUS | | | |
| UNKNOWN | 2778 | 164 | 5.6 |
| HER2+ | 156 | 6 | 3.7 |
| HR-/HER2- | 150 | 6 | 3.8 |
| HR+/HER2- | 522 | 21 | 3.9 |
| NA | 165 | 1 | 0.6 |
| TUMOR HISTOLOGY | | | |
| IDC | 2608 | 109 | 4.0 |
| ILC | 449 | 43 | 8.7 |
| MIXED | 136 | 10 | 6.8 |
| OTHERS | 538 | 36 | 6.3 |
| UNKNOWN | 40 | 0 | 0.0 |
| TOTAL PATIENTS | 3969 | 198 | 5.0 |

Abbreviations: ILC: invasive lobular carcinoma, IDC: invasive ductal carcinoma, HR: hormone receptor, NA: not available.

**Supplementary Table 2**. PIK3CA alterations counts in different gene domains for hypermutated samples with dominant APOBEC activity and hypermutated tumors without dominant APOBEC activity.

| Protein Change | Number of mutations in hypermutated tumors with dominant APOBEC | Number of mutations in hypermutated tumors without dominant APOBEC | *PIK3CA* Domains |
|---|---|---|---|
| p.C378F | 2 | 0 | C2 |
| p.C420R | 0 | 1 | C2 |
| p.D603H | 1 | 0 | Helical |
| p.E1012Q | 1 | 0 | Kinase |
| p.E365K | 1 | 1 | C2 |
| p.E418K | 0 | 2 | C2 |
| p.E453_L456del | 0 | 1 | C2 |
| p.E453K | 3 | 0 | C2 |
| p.E542K | 17 | 4 | Helical |
| p.E542Q | 1 | 0 | Helical |
| p.E545A | 1 | 0 | Helical |
| p.E545K | 30 | 5 | Helical |
| p.E545R | 2 | 0 | Helical |
| p.E726K | 11 | 1 | NA |
| p.E81K | 4 | 1 | ABD |
| p.G1049R | 2 | 0 | Kinase |
| p.G106V | 0 | 2 | NA |
| p.G118D | 0 | 2 | NA |
| p.H1047L | 2 | 1 | Kinase |
| p.H1047R | 23 | 12 | Kinase |
| p.H450_D454del | 1 | 0 | C2 |
| p.K240Q | 1 | 0 | RBD |
| p.L456L | 1 | 0 | C2 |
| p.L989V | 1 | 0 | Kinase |

| | | | |
|---|---|---|---|
| p.M1004I | 1 | 0 | Kinase |
| p.N145N | 0 | 1 | NA |
| p.N345K | 3 | 1 | C2 |
| p.N457_V461del | 0 | 1 | C2 |
| p.N457K | 1 | 0 | C2 |
| p.P366R | 0 | 1 | C2 |
| p.Q546H | 0 | 1 | Helical |
| p.Q546K | 1 | 0 | Helical |
| p.Q546P | 0 | 1 | Helical |
| p.Q75E | 0 | 1 | ABD |
| p.R108_I112delinsV | 0 | 1 | NA |
| p.R992* | 0 | 1 | Kinase |
| p.T1025A | 1 | 0 | Kinase |
| p.Y1021C | 0 | 2 | Kinase |

Abbreviations: APOBEC: Apolipoprotein B mRNA-editing enzyme catalytic polypeptide-like 3

**Supplementary Table 3** - Clinical-pathological characteristics and therapies records of patients with hypermutated breast cancer treated with pembrolizumab-based therapies.

| | Patient 1 | Patient 2 | Patient 3 |
|---|---|---|---|
| **Sex, age at metastatic disease diagnosis** | Female, 57 years-old | Female, 62 years-old | Female, 60 years-old |
| **Pathological diagnosis at time of metastatic recurrence** | Invasive carcinoma ER-/PR-/HER2- | Invasive carcinoma ER+/PR-/HER2- | Invasive carcinoma ER-/PR-/HER2- |
| **Prior systemic treatments in the adjuvant setting** | 1. AC followed by adjuvant ovarian suppression plus Anastrozole | 1. Adjuvant CMF followed by adjuvant Tamoxifen<br>2. Neoadjuvant TC and "adjuvant" anastrozole due to local recurrence | 1. Adjuvant CAF (right sided breast cancer)<br>2. Adjuvant CMF (second primary in the contralateral breast)<br>3. Adjuvant clinical trial (new primary in the right breast) |
| **Prior systemic treatments in the metastatic setting** | 1. Letrozole plus Palbociclib<br>2. Clinical trial | 1. Fulvestrant + Capecitabine<br>2. Exemestane + Everolimus<br>3. Clinical trial<br>4. Eribulin<br>5. Liposomal doxorubicin<br>6. Letrozol plus Palbociclib<br>7. Carboplatin<br>8. Gemcitabine | 1. Capecitabine<br>2. Navelbine<br>3. Paclitaxel<br>4. Eribulin<br>5. Cisplatin<br>6. Clinical trial |
| **Immunotherapy regimen received** | Pembrolizumab plus Eribulin | Pembrolizumab plus Nab-Paclitaxel | Pembrolizumab |
| **Type of objective response** | Complete response | Partial response | Partial response |

Abbreviations: AC: doxorubicin plus cyclophosphamide; CAF: cyclophosphamide, doxorubicin plus 5-fluorouracil; CMF: cyclophosphamide, methotrexate plus 5-fluorouracil; ER: estrogen receptor; PR: progesterone receptor; TC: docetaxel plus cyclophosphamide

**Supplementary Figure 1.** TMB distribution across clinical and pathological subgroups for whole breast cancer cohort. (A) Correlation between TMB (y-axis, log scale) and age at diagnosis (B) Boxplot indicating the mutation burden across IDC and ILC histology subtypes. Difference between the median TMB is not significant. (C) Boxplot indicating significant difference in median TMB across receptor subtypes. Numbers in parenthesis represents the total number of patients included in this analysis for each subgroup.

Abbreviations: ILC: invasive lobular carcinoma, IDC: invasive ductal carcinoma, TMB: tumor mutational burden

### Supplementary Figure 1

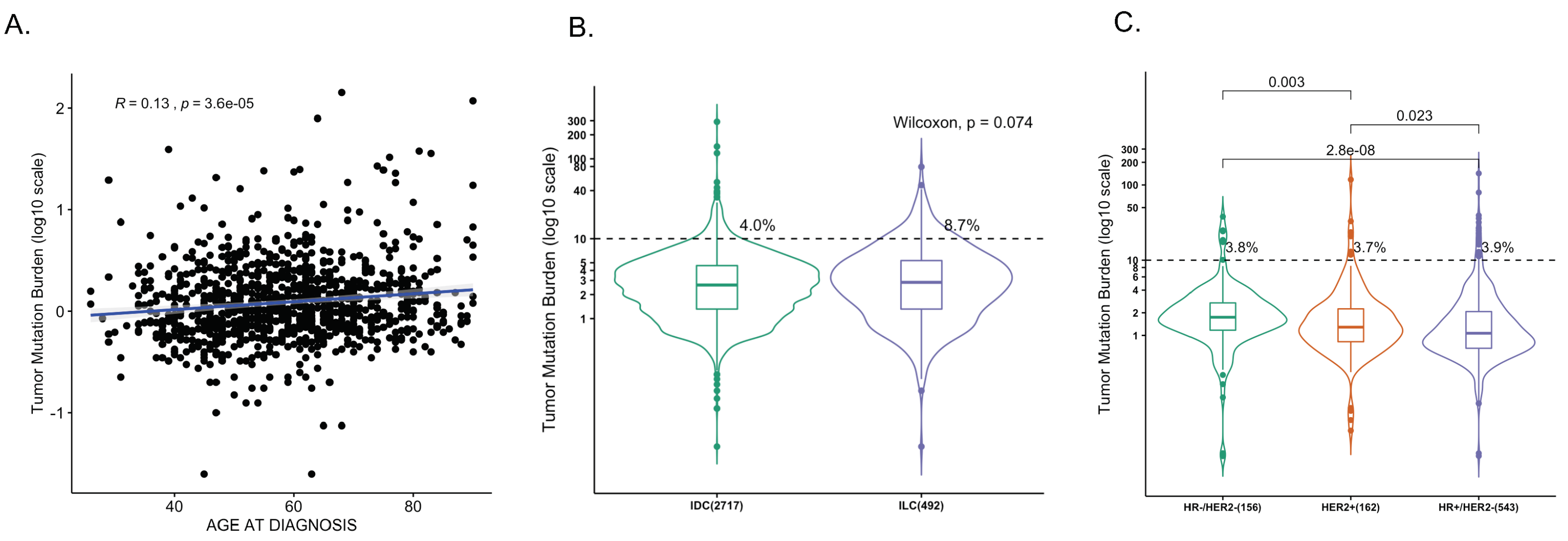

**Supplementary Figure 2.** Hypermutated breast cancers have a higher cytolytic score and higher neoantigen burden (a) Cytolytic activity is shown for tumors from the TCGA-BRCA cohort. Cytolytic activity is significantly higher for hypermutated tumors (TMB ≥10) vs normal tumors (TMB < 10). (b) Neoantigen burden is shown for tumors from the MBCProject cohort. Neoantigen burden is significantly higher for hypermutated tumors (TMB ≥10) versus normal tumors (TMB < 10). P-values are calculated using Wilcoxon rank-sum test. Numbers in parenthesis represents the total number of samples included in this analysis for each subgroup
Abbreviations: TMB: tumor mutational burden

Supplementary Figure 2

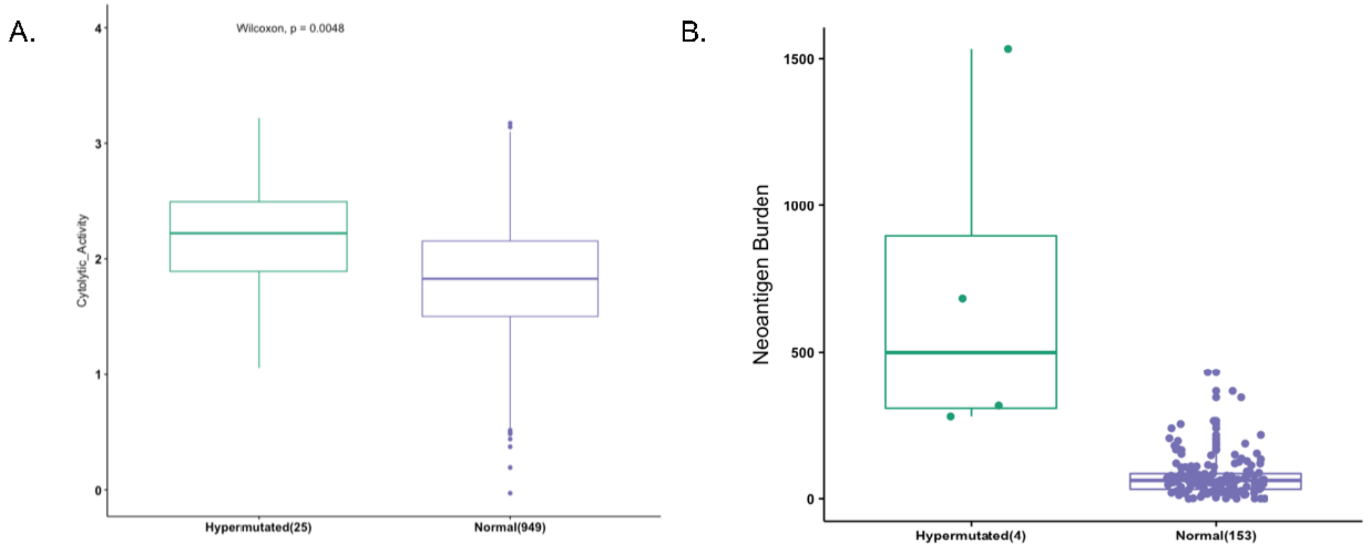
